# Group decision-making is optimal in adolescence

**DOI:** 10.1101/412726

**Authors:** Simone PW Haller, DAN Bang, Bahador Bahrami, Jennifer YF Lau

## Abstract

Group decision-making is required in early life in educational settings and central to a well-functioning society. However, there is little research on group decision-making in adolescence, despite the significant neuro-cognitive changes during this period. Researchers have studied adolescent decision-making in ‘static’ social contexts, such as risk-taking in the presence of peers, and largely deemed adolescent decision-making ‘sub-optimal.’ It is not clear whether these findings generalise to more dynamic social contexts, such as the discussions required to reach a group decision. Here we test the optimality of group decision-making at different stages of adolescence. Pairs of male pre-to-early adolescents (8 to 13 years of age) and mid-to-late adolescents (14 to 17 years of age) together performed a low-level, perceptual decision-making task. Whenever their individual decisions differed, they were required to negotiate a joint decision. While there were developmental differences in individual performance, the joint performance of both adolescent groups was at adult levels (data obtained from a previous study). Both adolescent groups achieved a level of joint performance expected under optimal integration of their individual information into a joint decision. Young adolescents’ joint, but not individual, performance deteriorated over time. The results are consistent with recent findings attesting to the competencies, rather than the shortcomings, of adolescent social behaviour.

## Introduction

Adolescence, spanning the second decade of life, is a period of profound physical and psychological development. It is characterised by continued brain maturation through synaptic pruning and myelination^1,2^, in particular of the prefrontal cortex, which is involved in higher-order functions such as decision-making, cognitive control and social cognition^3,4,5,6^. Indeed, the protracted development of prefrontal cortex has been linked to a range of ‘sub-optimal’ behaviours in adolescence, such as impaired monitoring of task performance and excessive risk-taking, especially in social contexts^4^. Surprisingly, there is little research on how the profound neuro-cognitive changes associated with adolescence affect the ability of adolescents to make decisions as part of a group. Group decision-making is required in early life in educational settings and, more generally, is central to a functioning society.

Here we report on a study which tested the optimality of group decision-making at different stages of adolescence using a well-established psychophysical paradigm^7^. While the choice problem in this paradigm is simple, the task captures important aspects of real-world social interaction: participants must resolve a disagreement with a same-aged friend through free, unconstrained verbal communication. Disagreements with peers over uncertain information play a central role in adolescents’ everyday life^8^. In addition, integrating and weighing social information is an important developmental task in adolescence, especially in the context of conflict resolution^9,10^.

### Decision-making and cognition in adolescence

Research on social decision-making in adolescence has largely pursued three avenues: i) individual learning in social contexts (i.e., “collaborative learning”; e.g.,^11,12^), ii) decision-making, and in particular risk-taking, in the presence of peers (e.g.,^4,13,14,15^); and iii) social negotiations and exchanges, often in the context of economic games (e.g.,^16,17,18^).

Research on collaborative learning has largely focused on educational contexts and individual learning outcomes (e.g.,^19,12^). There is converging evidence that working as part of a group can boost individual learning^20,21^, with key factors being exchange of ideas through communication^22,23^, the ability of other group members^24^ and the type of task^25^. However, it remains unknown whether these individual benefits translate into benefits for the group as a whole.

Research on social decision-making has largely been concerned with peer influence. Peers have a particularly salient, motivating effect on adolescent behaviour compared to other age groups. For example, peers are thought to increase adolescents’ proneness to engage in risky behaviours (e.g., risky behaviour in driving simulation games^15^). Recent studies have extended this work to suggest that the direction of these effects depend on the characteristics of the peer^26,27,28^ and the type of risk context (i.e., ambiguous vs known risk^29^). It has been proposed that increased susceptibility to peer influence is at least in part adaptive, resulting in important exploratory behaviour and formation of social networks outside of the family^29,30^. However, in the context of group decision-making, such increased susceptibility to peer influence may prevent a group from assigning appropriate weights to the opinions of its members^31^.

Research on social negotiations and exchanges has largely been concerned with economic behaviours. For instance, Guroglu et al.^32^ found that investment decisions in 8- to 18-year-olds in a version of the Ultimatum Game were increasingly modulated by the perceived tendency of another peer to reject a selfish offer. Burnett-Heyes et al.^16^ found developmental differences between mid- and late-adolescents in resource allocation: older adolescents were more likely to consider peers’ reciprocated feelings of friendship when deciding how to allocate resources in a modified Dictator Game. In the context of group decision-making, the protracted development of fairness and reciprocity norms may result in overly egocentric behaviour and discounting of others’ opinions in adolescence.

In addition to social decision-making, there is also a body of research concerned with cognitive functions, which are relevant to decision-making and social interaction more generally. For instance, using the same basic perceptual task as we use here, Weil et al.^33^ found that the ability to accurately monitor and report on one’s task performance – an ability known as metacognition – continues to develop in adolescence and only levels off toward adulthood. Similarly, there is evidence that cognitive and affective aspects of social cognition continue to develop in adolescence^34,35^.

Overall, research on decision-making and cognition in adolescence indicates that, in certain domains such as high-risk domains, social context has unique effects on adolescent behaviour. Additionally, with increasing age, as social-cognitive skills are refined, social computations increase in sophistication, affecting how youth relate to peers. To our knowledge, no study has yet examined the optimality of group decision-making in adolescence. Critically, it is not clear whether the findings obtained in more ‘static’ social contexts, such as the manipulation of peer presence, extend to more ‘dynamic’ social contexts, such as the discussion preceding a group decision. Additionally, no study has used a paradigm that can isolate developmental differences at the group level over and above developmental differences at the individual level – crucial when testing hypothesis about group-level developmental effects.

### Current study

We used a psychophysical task to study group decision-making in male pre-to-early adolescents (8- to 13-year-olds) and mid-to-late adolescents (14- to 17-year-olds). On each trial, pairs of participants viewed two brief displays serially, one of them containing a faint target. Participants privately indicated which display they thought contained the target. In the case of agreement (i.e., they privately selected the same display), they received feedback about choice accuracy and continued to the next trial. In the case of disagreement (i.e., they privately selected different displays), they were first required to reach a group decision. In contrast to commonly used social tasks based on abstract social scenarios (e.g., economic games or interactions with virtual agents), the paradigm involves free social interaction with a close peer and closely resembles a common type of everyday social experience, such as when two friends discuss whether a ball crossed the line in a football match. As the paradigm allows for standardisation of joint performance by individual performance, it is particularly suitable for studying group decision-making across development.

Bahrami et al.^7^ found that a Weighted Confidence Sharing (WCS) model provided the best fit to adult data collected on the above task. According to the WCS model, to resolve disagreement, participants communicate the level of confidence in their respective opinions and then weight each opinion by the communicated confidence. The model assumes that participants can accurately estimate the reliability of their private opinion and accurately communicate this information – a set of skills encompassing metacognition^36^ and social cognition^37,38^ – and thus provides an upper bound on joint performance. Critically, the ‘optimality’ measure derived from the WCS model takes into account individual performance and thus provides a measure of joint performance that is unbiased by low-level developmental differences. What do we expect in adolescence? Given the protracted development of decision-making, social cognition and metacognition, we predicted joint performance to be lower and less optimal in the younger than in the older adolescent group – and that neither age group would reach an adult level of joint performance as estimated from the dataset collected by Bahrami et al.^7^.

## Methods

### Sample

We recruited seventy-four adolescent participants in pairs (37 dyads in total) via adverts in the local community. The members of each dyad were friends with one another. We recruited two age groups: 40 pre-to-early adolescents (20 dyads; mean age ± SD = 11.0 ± 1.11; age range: 8.5-12.6 years; school years: 5-8) and 34 mid-to-late adolescents (17 dyads; mean age ± SD =15.5 ± .79; age range: 14.1-16.8 years; school years 9-11). All participants were healthy males (for consistency with Bahrami et al.,^7^) with normal or corrected-to-normal vision. Participants were paid £20 for participation. We obtained written assent from participants and written informed consent from legal guardians. The University of Oxford Central University Research Ethics Committee (CUREC) approved the study (Title: Collective decision-making, Protocol Number: MSD/IDREC/C1/2012/16). The study was performed in accordance with all relevant guidelines and regulations. An adult dataset (14 dyads; mean age ± SD = 29.6 ± 7.7; age range: 18.3-50.2 years) was obtained from Experiment 1 in Bahrami et al.^7^ where participants also were recruited in pairs of friends.

### Task

Participants performed a two-interval forced-choice contrast-discrimination task as part of a dyad (see **Figure 1** for a schematic of an experimental trial). Participants sat at right angles to each other in a dark room, with their own monitor and response device (keyboard or mouse). On each trial, participants were presented with two consecutive viewing displays, each containing six vertically oriented Gabor patches. In one of the two displays, the contrast level of one of the six Gabor patches (the target) was increased by adding one of four values (.075,.15, .20, .30) to its baseline contrast (.15). We chose the values for the current study based on four pilot adolescent participants. The values differ from those used for the adult data (baseline contrast: .10; added contrast: .015, .035, .07, .15), but we note that our key measures of joint performance are normalised relative to individual performance and thus comparable across datasets. Target location and display was randomized across trials and experimental sessions, contrast values were counterbalanced across trials such that each appeared equal number of times. Participants viewed identical visual stimuli on each trial.

**Figure 1.**
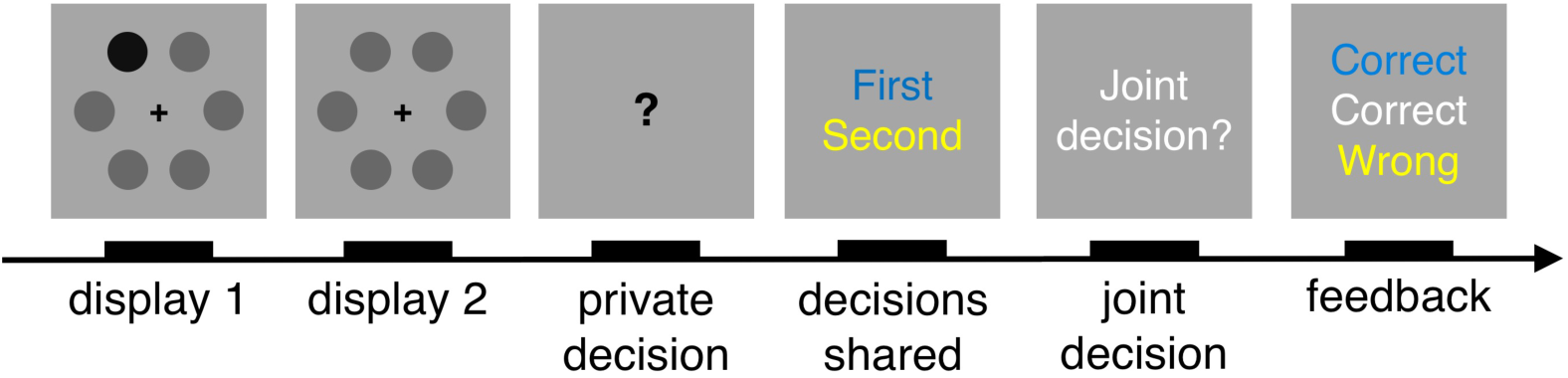
Schematic of experimental procedure. On each trial, participants viewed two consecutive displays, each containing six contrast gratings (here shown as dots). There was a target with higher contrast (here the darker dot) in one of the two displays. Participants privately decided which display they thought contained the target. The private responses were shared. In the case of disagreement, participants were required to make a joint decision; they took turns at indicating the joint decision. Participants received feedback about the accuracy of the individual and joint decisions, before continuing to the next trial. In the case of agreement, participants proceeded immediately to feedback. Individual responses (blue and yellow) and joint responses (white) were identified by colours.

Trials were initiated by the participant with the keyboard after consulting with their partner. A black central fixation cross (width: 0.75 degrees visual angle) appeared on the screen for a variable period (500-1000 ms). After the stimulus presentation, separated by a blank display lasting 1000 ms, participants were asked to indicate which display they thought contained the target, without discussing their answers. A question mark prompt after the second display signalled to the participants to respond. The question mark remained on the screen until both participants had made a response. Once both of them had responded, the individual decisions were made public (keyboard: blue; mouse: yellow). In the case of agreement (i.e., they privately selected the same display), they received feedback (see below) and continued to the next trial. In the case of disagreement (i.e., they privately selected different displays), they were asked to agree on a joint decision through verbal discussion. Participants were free to discuss the joint decision as long as they wanted. Participants took turns at indicating the joint decision (keyboard: even trials; mouse: odd trials). Once the joint response had been made, they received feedback about the accuracy of each decision (keyboard: blue; mouse: yellow; joint: white) and continued to the next trial.

Participants first completed a practice block of 16 trials. They then performed two experimental sessions, each consisting of 8 blocks of 16 trials (128 trials in each session and 256 trials in total). The two sessions were separated by a short break (5-10 mins). Participants swapped response devices between the two sessions. See Bahrami et al.^7^ for details about display parameters, response mode and stimulus presentation.

### Procedure

We introduced the task as a picture-book game akin to *Where’s Wally*, replacing the Gabor patches with cartoon figures. As in the main task, in one of the two displays, one of the cartoon figures had a higher level of contrast. We ensured that participants had understood the basic premise of the task (i.e., to respond whether the target was in the first or the second display), before introducing the Gabor patches. Participants were told that the task was about teamwork and they were encouraged to try to make as many correct joint decisions as possible. An experimenter was present throughout the entire study to ensure that all instructions were observed. Parents were not present during the task. The study lasted about two hours including breaks.

### Measures

*Individual performance*

Accuracy

We calculated accuracy as fraction of correct individual decisions.

Reaction time

Reaction time was calculated as seconds taken to make a decision.

### Sensitivity

To estimate sensitivity for each dyad member, we first plotted the proportion of trials on which the target was reported to be in the second display against the difference in contrast between the second and the first display at the target location. The data points were then fit with a cumulative Gaussian function whose parameters were bias, *b*, and variance, *σ*^2^ – the parameters were estimated using a probit regression model as implemented by MATLAB’s (Mathworks Inc.) *glmfit* function. A participant with bias *b* and variance *σ*^2^ would have a psychometric curve, denoted *p*(Δ*c*), where Δ*c* is the contrast difference between the second and first display at the target location, given by

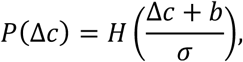

where *H(Z)* is the cumulative normal function

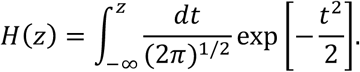

The psychometric curve, *p*(Δ*c*), tells us how the probability of reporting that the target is in the second display changes with contrast difference. Given the above definition, the variance in responses is related to the maximum slope of the psychometric curve, denoted *S*, via

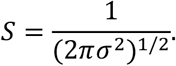

A steep slope indicates small variance and thus highly sensitive performance. In contrast to accuracy, sensitivity provides a bias-free measure of performance.

### Joint performance

#### Joint accuracy

We calculated joint accuracy as fraction of correct joint decisions (both agreement and disagreement trials).

#### Joint reaction time

We calculated joint reaction time as seconds taken to reach a joint decision.

#### Egocentric bias

To quantify ‘egocentric bias’, we computed the proportion of joint-decision trials in which a dyad member indicated the joint decision and the joint decision was the same as that made by the dyad member^39^.

#### Similarity

We computed the similarity of dyad members’ sensitivities as the ratio of the sensitivity of the worse dyad member to that of the better dyad member, *S*_min_/*S*_max_, with values near zero corresponding to dyad members with very different sensitivities and values near one corresponding to dyad members of nearly equal sensitivity.

#### Joint sensitivity

Joint sensitivity was quantified using the same procedure as for individual sensitivity but this time relating joint responses to the stimulus.

#### Collective benefit

We computed the collective benefit accrued by a dyad as the ratio of the sensitivity of the dyad to that of the more sensitive dyad member, *S*_dyad_/*S*_max_, with values below 1 indicating a collective loss and values above 1 indicating a collective benefit.

#### Optimality

We estimated the ‘optimality’ of joint performance using the Weighted Confidence Sharing (WCS) model developed by Bahrami et al.^7^. The model assumes that participants on each trial can accurately estimate and communicate the reliability of their respective individual decisions– with the joint decision reached by weighting the individual decisions by the communicated reliabilities. The upper boundary on joint performance can thus be defined as

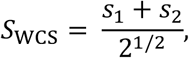

where *s*_1_ and *s*_2_ are the individual sensitivities calculated as above. We computed our optimality index as the ratio of the dyad’s sensitivity to that predicted by the WCS model, *S*_dyad_/*S*_wcs_, with values above 1 indicating that the dyad is ‘supra-optimal’ and values below 1 indicating that the dyad is ‘sub-optimal’. See Bahrami et al.^7^ for mathematical details.

### Data exclusion

We excluded dyads where one or both of the members performed at 55% accuracy or lower and/or had a negative sensitivity (18 pre-to-early adolescent and 16 mid-to-late adolescent dyads remaining after data exclusion).

## Results

### Individual performance

#### Accuracy

The mid-to-late adolescents (OA: older adolescents) reached higher levels of accuracy than the pre-to-early adolescents (YA: younger adolescents) (OAs: mean ± SD = .777 ± .074; YAs: mean ± SD = .701 ± .077) (**Figure 2A**; *t*(66) = 3.745, *p* < .001, *d* = .930, independent-samples *t*-test). YAs, however, reached acceptable levels of performance (about 70%), indicating that the task was not too difficult. We emphasise that our main measures of joint performance are normalised by individual performance and therefore can be compared between age groups despite differences at the individual level.

**Figure 2.**
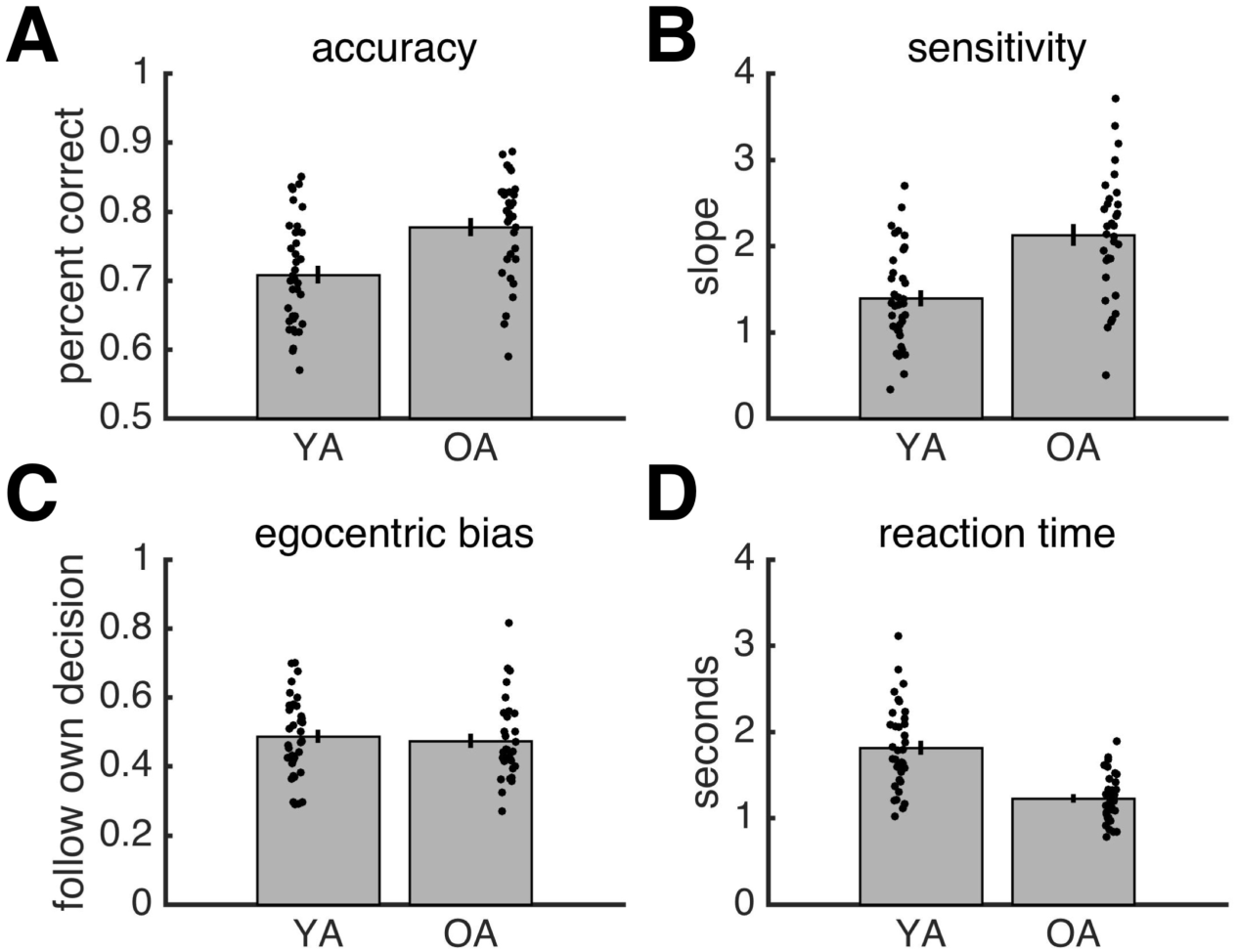
Results for individual behaviour. **A**, Accuracy. **B**, Sensitivity. **C**, Egocentric bias. **D**, Reaction time. **A-D**, Grey bars are data averaged across individuals. Error bars reflect SEM. Each black dot is an individual. YA: younger adolescents. OA: older adolescents.

#### Sensitivity

OAs reached higher levels of sensitivity (i.e., steepness of psychometric function) than YAs (OAs: mean ± SD = 2.128 ± .716; YAs: mean ± SD = 1.394 ± .561) (**Figure 2B**; *t*(66) = 4.731,

*p* < .001, *d* = 1.165, independent-samples *t*-test).

#### Egocentric bias

Both YAs (mean ± SD = 0.487 ± .115) and the OAs (mean ± SD = 0.474 ± .117) opted for their own choice about half of the time. There was no significant difference in egocentric bias between the age groups (**Figure 2C**; *t*(66) = 0.469, *p* = .641, *d* = .115, independent-samples *t*-test).

#### Reaction time

OAs made faster responses than YAs (OAs: mean ± SD = 1.227 ± .285 seconds; YAs: mean ± SD = 1.814 ± .488) (**Figure 2D**; *t*(66) = -5.960, p < .001, *d* = -1.467, independent-samples *t*-test).

### Joint performance

#### Accuracy

As expected given the differences in individual performance, OA dyads (mean ± SD = .848 ± .060) reached higher levels of accuracy than YA dyads (mean ± SD = .755 ± .076) (*t*(32) = 3.897, *p* < .001, *d* = 1.378, independent samples *t*-test).

#### Similarity

The similarity of dyad members’ sensitivities was on average lower in the YA group (mean ± SD = .698 ± .191) than in the OA group (mean ± SD = .755 ± .230). However, due to the within-group heterogeneity, the difference in similarity was not significant (**Figure 3A**; *t*(32) = -0.793, *p* = .433, *d* = -.280, independent-samples *t*-test).

**Figure 3.**
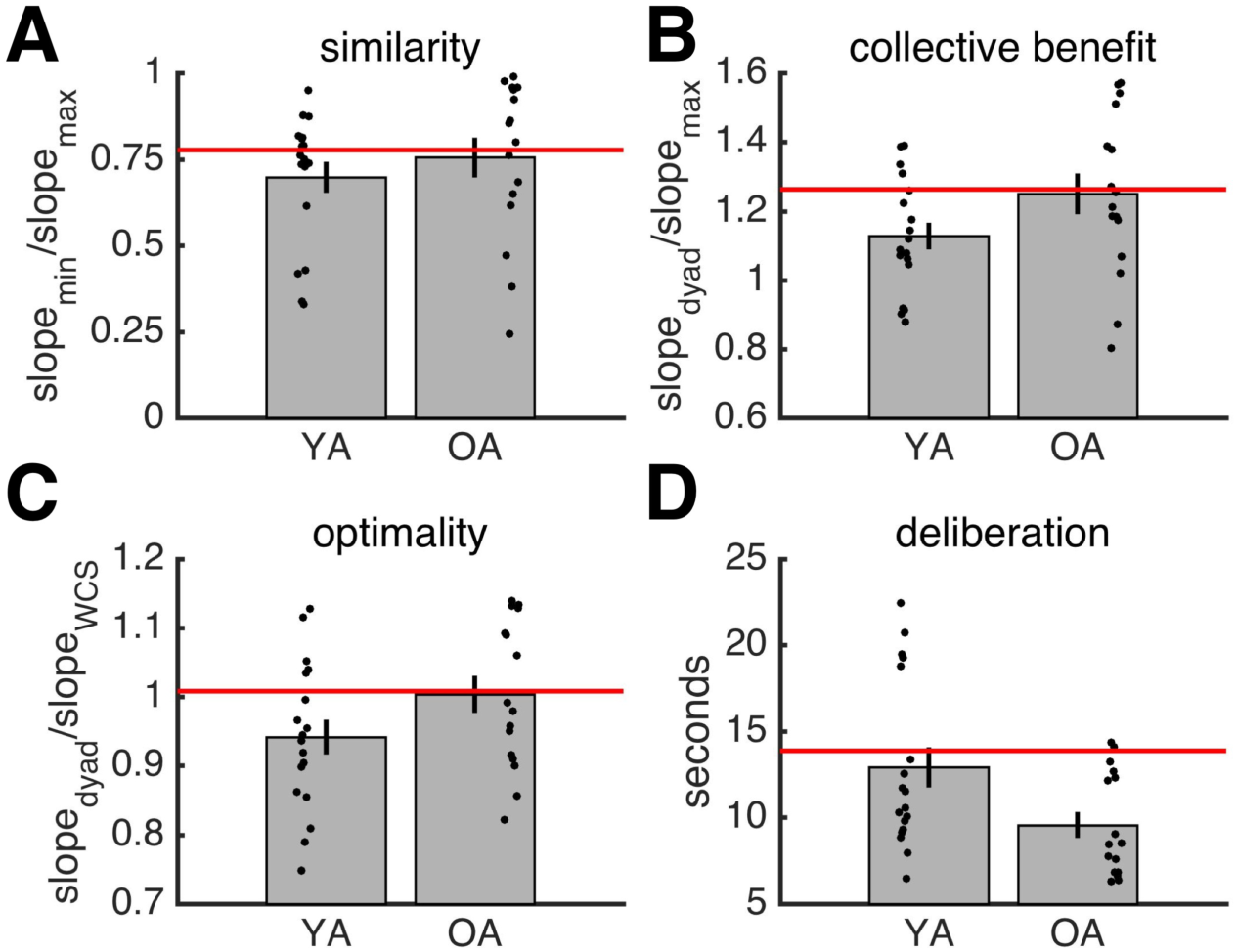
Results for joint behaviour. **A**, Similarity of sensitivity. **B**, Collective benefit. **C**, Optimality. **D**, Deliberation time. **A-D**, Horizontal red lines indicate mean of adult data. Grey bars are data averaged across individuals. Error bars reflect SEM. Each black dot is a dyad. YA: younger adolescents. OA: older adolescents.

#### Similarity

The similarity of dyad members’ sensitivities was on average lower in the YA group (mean ± SD = .698 ± .191) than in the OA group (mean ± SD = .755 ± .230). However, due to the within-group heterogeneity, the difference in similarity was not significant (**Figure 3A**; *t*(32) = -0.793, *p* = .433, *d* = -.280, independent-samples *t*-test).

**Figure 3.**
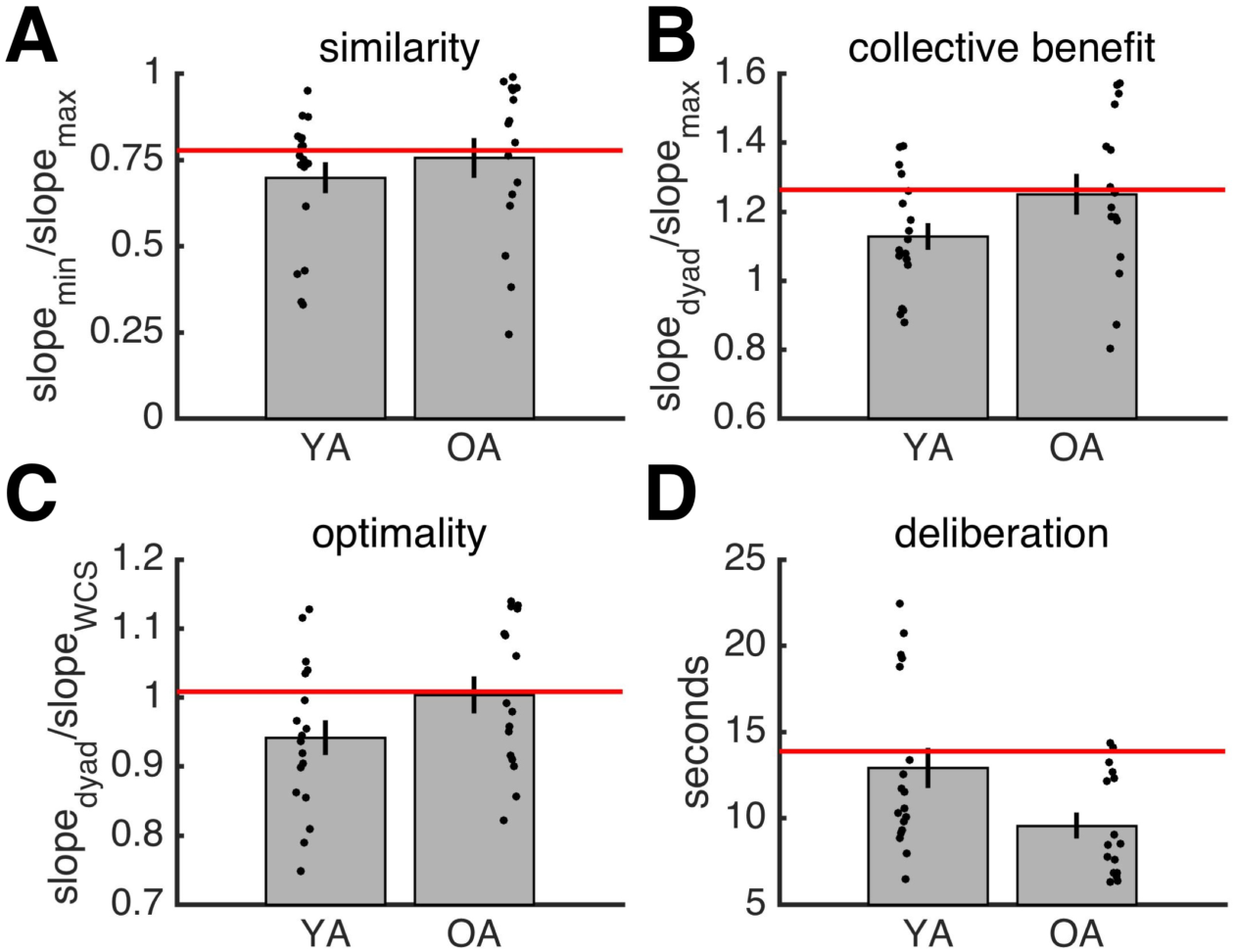
Results for joint behaviour. **A**, Similarity of sensitivity. **B**, Collective benefit. **C**, Optimality. **D**, Deliberation time. **A-D**, Horizontal red lines indicate mean of adult data. Grey bars are data averaged across individuals. Error bars reflect SEM. Each black dot is a dyad. YA: younger adolescents. OA: older adolescents.

#### Joint sensitivity

OA dyads (mean ± SD = 3.014 ± .878) reached higher levels of sensitivity than YA dyads (mean ± SD = 1.832 ± .558) (*t*(32) = 12.134, *p* < .001; *d* = 4.290, independent-samples *t*-test).

#### Collective benefit

YA and OA dyads on average outperformed the more sensitive dyad members (*S*_HIFH_/*S*_CFG_, **Figure 3B**; YA: *t*(17) = 3.288, *p* = .004; OA: *t*(15) = 4.233, *p* < .001; one-sample *t*-test, null model mean is 1). The collective benefit was on average higher in the OA group (mean ± SD = 1.250 ± .236) than in the YA group (mean ± SD = 1.128 ± .165). However, this difference was not significant (**Figure 3B**; *t*(32) = -1.767, *p* = .087, *d* = -.625, independent-samples *t*-test).

#### Optimality

Comparison of the empirically observed joint sensitivity to that expected under optimal performance showed that YA dyads on average were ‘sub-optimal’ whereas OA dyads on average were ‘optimal’ (**Figure 3C**; YA: *t*(17) = -2.321, *p* = .033; OA: *t*(15) = 0.127, *p* = .901; one-sample *t*-test, null model mean is 1). In line with the results for collective benefit, while OA dyads achieved a higher level of optimality (mean ± SD = 1.003 ± .108) than YA dyads (mean ± SD = 0.942 ± .107), this difference was not significant (**Figure 3C**; *t*(32) = -1.681, *p*= .103, *d* = -.594, independent-samples *t*-test).

#### Reaction time

YA dyads (mean ± SD = 12.901 ± 4.928 secs) took slightly but significantly longer than OA dyads (mean ± SD = 9.564 ± 3.000 secs) to reach joint decisions (**Figure 3D**; *t*(32) = 2.348, *p* = .025, *d* = .830, independent-samples *t*-test).

#### Adult benchmark

Direct group-comparisons using the optimality index showed no differences between, on the one hand, YA dyads and OA dyads and, on the other hand, adult dyads (adults: mean ± SD = 1.008 ± 0.1432) (**Figure 3C**; YA vs. adults: *t*(32) = -1.506, *p* = .143, *d* = -.550; OA vs. adults: *t*(30) = -0.100, *p* = .921; *d* = -.035, independent-samples *t*-test). Hence, the three age groups reached comparable levels of joint performance given their individual performance. In other words, no developmental difference emerged when individual performance was controlled for.

### Session analyses

To test for group differences in the temporal profile of individual and joint behaviour, we split the data into two sessions (participants performed two sets of 128 trials divided by a short break), and applied repeated-measures ANOVAs with session as within-subject factor and age group as between-subject factor to our measures of individual and joint performance. No age group by session interaction emerged for individual or joint behaviour (**Figure 4;** individual behaviour: accuracy: *F*_1,66_ = 0.047, *p* = .829; sensitivity: *F*_1,66_ = 0.060, *p* = .807; **Figure 5;** joint behaviour: collective benefit *F*_1,32_ = 2. 458, *p* = .127; optimality *F*_1,32_ = 2.037, *p* = .163).

**Figure 5.**
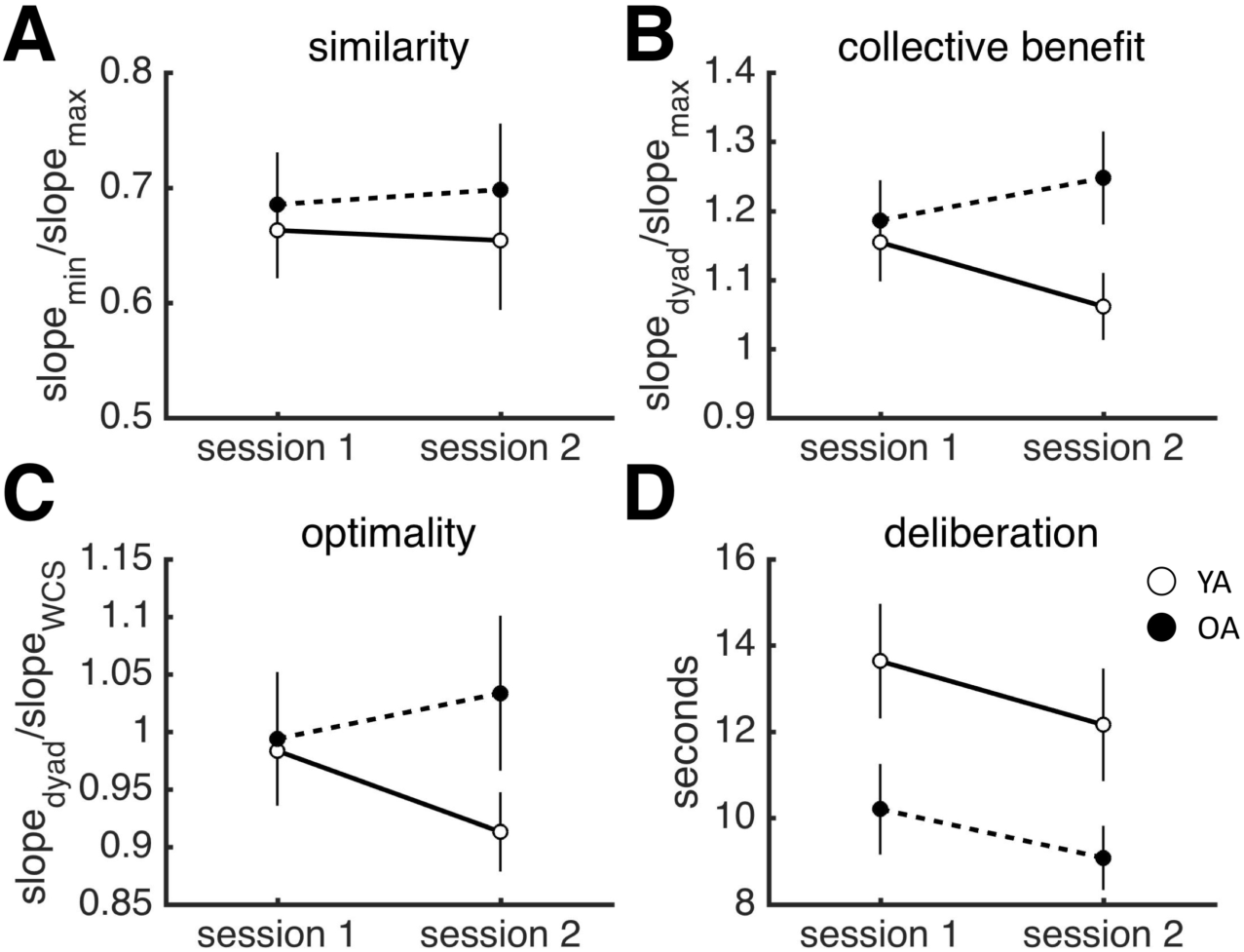
Results for joint behaviour by session. **A**, Similarity of sensitivity. **B**, Collective benefit. **C**, Optimality. **D**, Deliberation time. Dots are data averaged across dyads. Error bars reflect SEM. YA: younger adolescents. OA: older adolescents.

**Figure 4.**
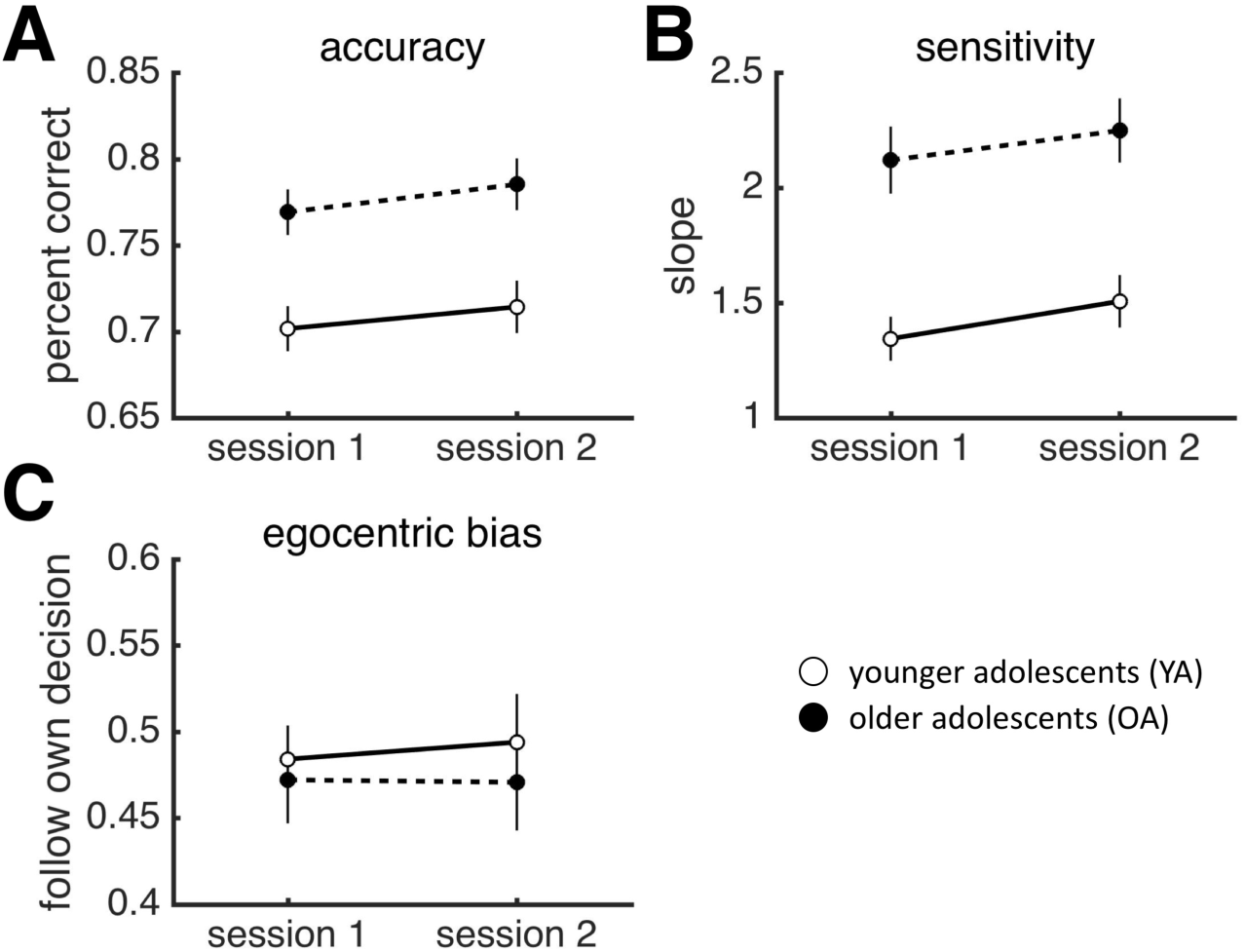
Results for individual behaviour by session. **A**, Similarity of sensitivity. **B**, Collective benefit. **C**, Optimality. **D**, Deliberation time. Dots are data averaged across individuals. Error bars reflect SEM. YA: younger adolescents. OA: older adolescents.

However, when examined per session, YA did not reap a collective benefit during the second session while OA outperformed the more sensitive member of the group in both sessions (YA: Session 1: *t*(17) = 2.707, *p* = .014; Session 2: *t*(17) = 1.269, *p* = .222; OA: Session1: *t*(15) = 3.208, *p* = .006; Session 2: *t*(15) = 3.679, *p* = .002; one-sample *t*-test, null model mean is 1). Additionally, YA dyads were ‘sub-optimal’ only during the second session – YA dyads were, in fact, ‘optimal’ during the first session, while OA remained ‘optimal’ across both sessions (Session 1: YA: *t*(17) = -0.383, *p* = .707; Session 2: *t*(17) = -2.533, *p* = .021; OA: *t*(15) = -0.167, *p* = .686; Session 2: *t*(15) = 0.850, *p* = .401; one-sample *t*-test, null model mean is 1). Hence, any developmental differences in the aggregate data are likely the result of YA’s performance during the second session, where YA performed worse as a group, but not individually. A direct comparison of the measures of joint performance separately for each session identified differences in the second session only (Session 1: Collective benefit: *t*(32) = 0.393, *p* = .697; Optimality: *t*(32) = 0.181, *p* = .858; Session 2: Collective benefit: *t(*32) = 2.275, *p* = .030; Optimality: *t*(32) = 2.317, *p* = .027).

### Continuous age

See *Supporting Information* for details on the age of each participant and additional analyses of individual and joint performance using age as a continuous variable. While the age range was wider in the younger age group (8-13) compared to the older age group (14-17), only three participants in the younger age group were below the age of 10 (**Figure S1**). Significant positive developmental gradients were found for individual performance (accuracy, sensitivity and reaction time, see **Figure S2**). Developmental gradients for joint performance were not significant (collective benefit and optimality, see **Figure S3**).

## Discussion

Group decision-making, integrating and weighing social information, is an important developmental milestone in adolescence. The aim of this study was to investigate the optimality of group decision-making in adolescence (male sample). We used a psychophysical task, which allowed us to precisely separate developmental differences in joint performance from developmental differences in individual performance. We examined two measures of joint performance: collective benefit (performance of the group relative to the group) and optimality (WCS model). We observed significant developmental differences in individual performance, with pre-to-early adolescents performing significantly worse than mid-to-late adolescents. Mid-to-late adolescents were significantly more accurate, more sensitive and faster to reach consensus than the younger age group. However, we found that, in terms of joint performance, both groups were able to accrue collective benefits and perform at optimal, adult levels. Interestingly, young adolescents’ joint performance deteriorated over time (while older adolescents’ showed performance gains over time), prompting one to think of fatigue and lapse of attention as the most likely cause of this temporal deterioration. This account, however, is inconsistent with the observation that the individual performance of young adolescents did *not* deteriorate over time. If anything, individuals improved over time in both groups (Fig 4A-B). A general loss of attention, drop in arousal or fatigue cannot explain the specific drop in collective performance across time in the younger adolescents (Fig 5B-C). These results suggest, perhaps surprisingly, that whereas social decision-making per se plateaus relatively early on in development, the ability to maintain social activities shows a more protracted development.

Research on adolescent decision-making in social contexts has largely focused on how peers increase the propensity of youth to make risky decisions^13,15^. This focus on immaturity or sub-optimality in adolescent decision-making has perhaps neglected that adolescents in fact display a range of sophisticated social behaviours outside the laboratory. In addition, studies examining both individual differences and age effects in cognitive performance have found that variance attributable to individual differences (e.g. variation in IQ within an age bracket) is often much larger than variance attributable to age. For instance, Roalf et al.^40^ found that, while performance on an N-back working memory task improved with age from 8 to 22 years, many late adolescents performed above the average young adult. In other words, age effects, while significant, may be relatively subtle in some domains. In order to build integrative and comprehensive developmental models of social decision-making, we need to use approaches which are not only naturalistic but also allow for precise quantification of behaviour and strategies.

The current findings can be the starting point for a line of research with direct implications for educational settings. While limited to simple perceptual decisions, the current findings indicate that even individuals as young as ten years of age can benefit from social interaction to improve joint performance. Future research should examine whether the current findings translate to tasks used in educational settings, and examine whether benefits obtained from group decision-making can feed back into individual learning and improve individual learning outcomes.

The current study has several strengths. First, it adds to the limited literature on group decision-making in adolescence, which predominantly has been concerned with individual outcomes. In addition, it is important to extend our understanding of social decision-making in adolescence beyond the effects of peers on individual behaviour^41^. Second, the paradigm used in this study balances tight experimental control (i.e., precise and independent measurement of individual and joint performance) with ecological validity (i.e., naturalistic, unconstrained interaction in the setting of real-world friendships). Third, using a computational approach (i.e., the WCS model) is uniquely suited to further test hypotheses about specific decision-making strategies. In particular, the WCS model assumes that individuals in a dyad can accurately estimate and communicate the reliability of their decision on each trial – an effective strategy for individuals with relatively similar sensitivities. Using such an approach allows us to go beyond asking *whether* adolescents can achieve collective benefits to understand *how* adolescents are able to do so.

The current study also has its limitations. First, the rich but largely uncontrolled setting of the study (i.e., free discussion, real-world relationships) may have contributed to the absence of developmental effects. While a previous study in adults on a similar psychosocial task found no effect of familiarity on joint performance^42^, familiarity may play a role in efficient communication in social interactions in younger age groups. Future research should assess the effects of relationship type and communication format on joint performance in adolescents. It may be that unconstrained communication with a peer is key for younger individuals – with developmental effects becoming apparent when individuals are required to rely on abstract indicators to arrive at a joint decision (e.g., confidence ratings made on a scale) instead of freely communicating it. Second, the sample was entirely male. Because same sex friendships during adolescence may be qualitatively different in males and females^43^, it remains to be seen whether the current results extend to adolescent females. We note, however, that two adult studies using a similar task found comparable behavioural results across male-male, male-female and female-female groups^39,44^. Lastly, session-by-session analyses showed that joint performance declined in younger adolescents. Restricting trial number in future developmental studies may help make results more comparable, unless temporal aspects of performance are of primary interest.

While the task relies on unconstrained social interactions, other aspects of the task are constrained (i.e., stimuli, number of participants, two choices) in order to provide experimental control and allow for precise measurement and computational modelling. Future work may want to expand on the basic paradigm. Developmental effects in how information is integrated in group decisions may become apparent under increased social-cognitive load (i.e., different stimuli or multiple group members) or in a more affectively-laden context (i.e., dyads selected for specific interactive dynamics).

Adolescents are experts in their own social worlds, showing remarkable flexibility and abilities to adapt and learn in social contexts: they operate complex social networks and are often quick adopters of new social trends from fashion to music and social media^30^. Here we observed that group decision-making is largely optimal in adolescents, with even very young adolescents being able to perform at adult levels. These results underscore the importance of examining different aspects of social decision-making in adolescences and studying adolescent social interaction within the setting of authentic relationships.

## Acknowledgements

SH was supported by a studentship from Medical Research Council (Reference: 1242237) and a Scatcherd European Scholarship. DB was supported by the Calleva Research Centre for Evolution and Human Sciences. BB was supported by the European Research Council (NEUROCODEC 309865).

## Author Contributions

DB, BB and JL designed the study. DB collected the data. SH and DB analysed the data. SH, DB and JL wrote the first draft of the manuscript. All authors reviewed and approved the final draft of the manuscript.

## Data availability

Deidentified raw data and code supporting main analyses are available on GitHub (https://github.com/danbang/article-adolescence-optimal/).

## Supporting Information

### Age

**Figure S1.**
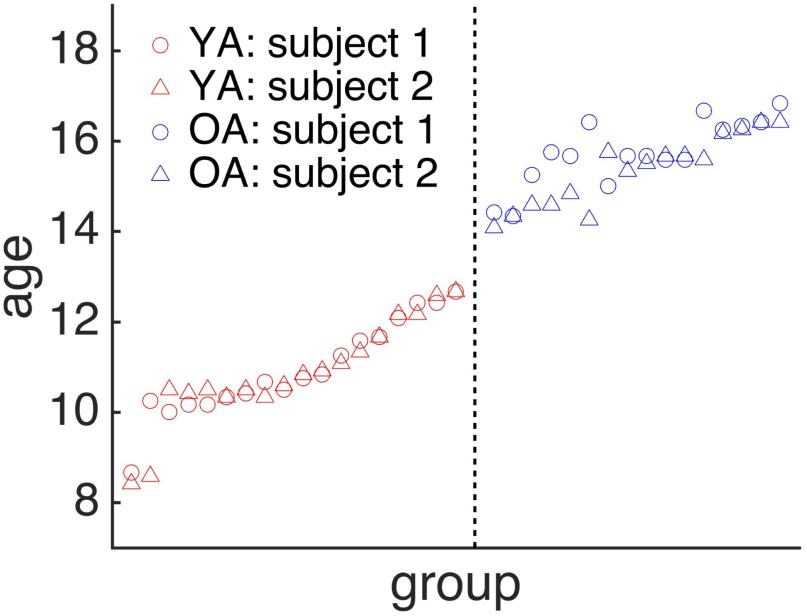
shows age in years for each dyad member in each age group. Age in years for all participants.

### Age as a continuous measure

To further investigate developmental differences in individual and joint behaviour, we used multiple linear regression to assess the relationship between age (individual age or the mean age of a dyad) and the measures of individual and joint performance. To take into account non-linear trajectories, we included both a linear term (age) and a quadratic term (age^2^) in our regression model.

#### Individual performance

We observed significantly positive developmental gradients for accuracy (**Figure S2A**), sensitivity (**Figure S2B**) and choice reaction time (**Figure S2C**). We did not observe a developmental gradient for egocentric bias (**Figure S2D**); while there were individual differences in egocentric bias (dot dispersion), these did not vary with age.

#### Joint performance

We did not observe a developmental gradient for similarity of sensitivity (**Figure S3A**). However, we observed marginally positive developmental gradients for collective benefit (**Figure S3B**) and optimality (**Figure S3C**) together with a marginally negative developmental gradient for deliberation time (**Figure S3D**). These results are most likely driven by the temporal deterioration of joint performance among younger participants (**Figure 5**).

**Figure S2.**
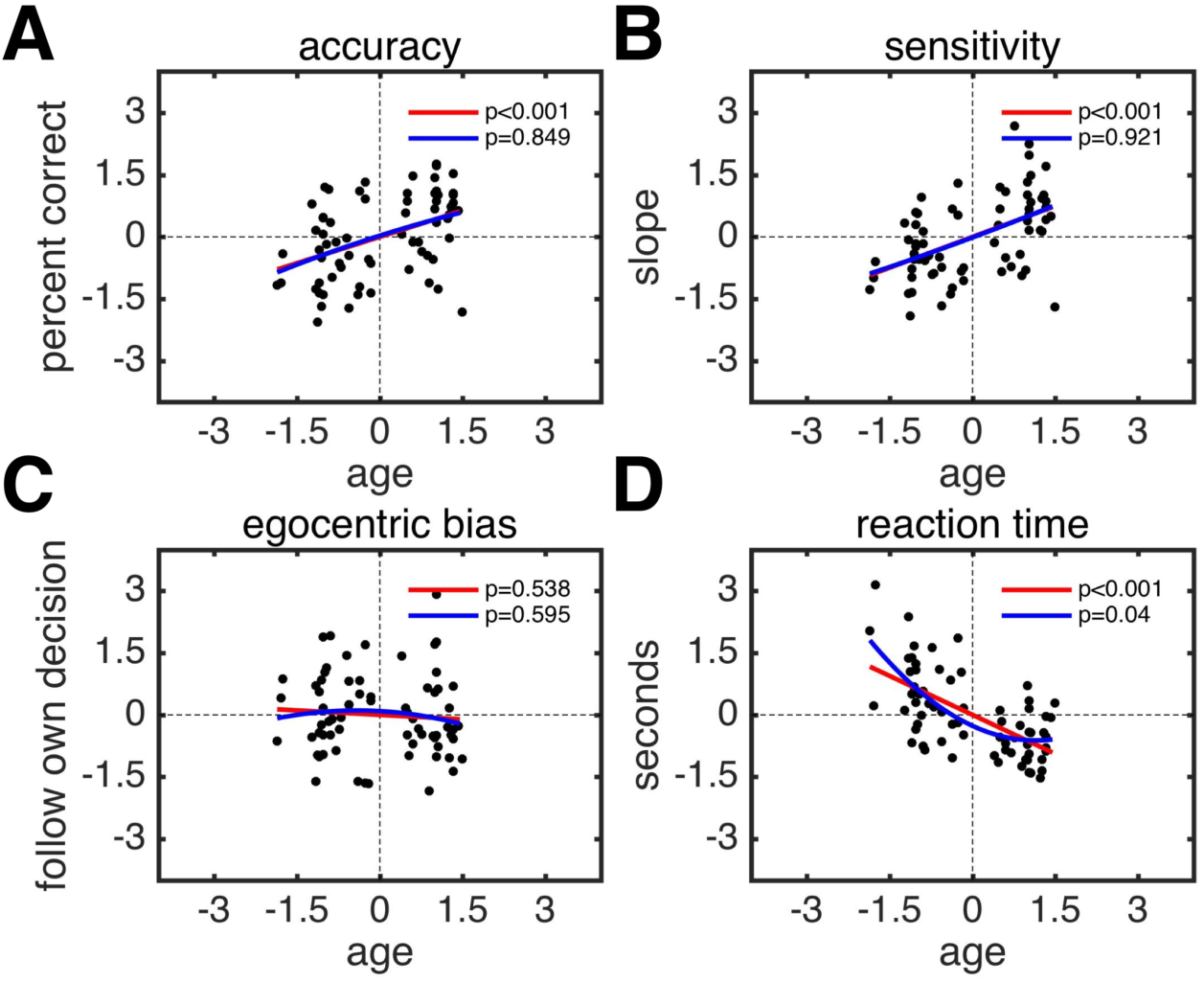
Developmental gradients for individual behaviour. **A**, Accuracy. **B**, Sensitivity. **C**, Egocentric bias. **D**, Reaction time. **A-D**, Each dot is a participant. The lines indicate the slopes for the linear (red) and the quadratic (blue) terms for age; the *p*-values indicate the significance of the terms. All variables were *z*-scored before being entered into the regression model.

**Figure S3.**
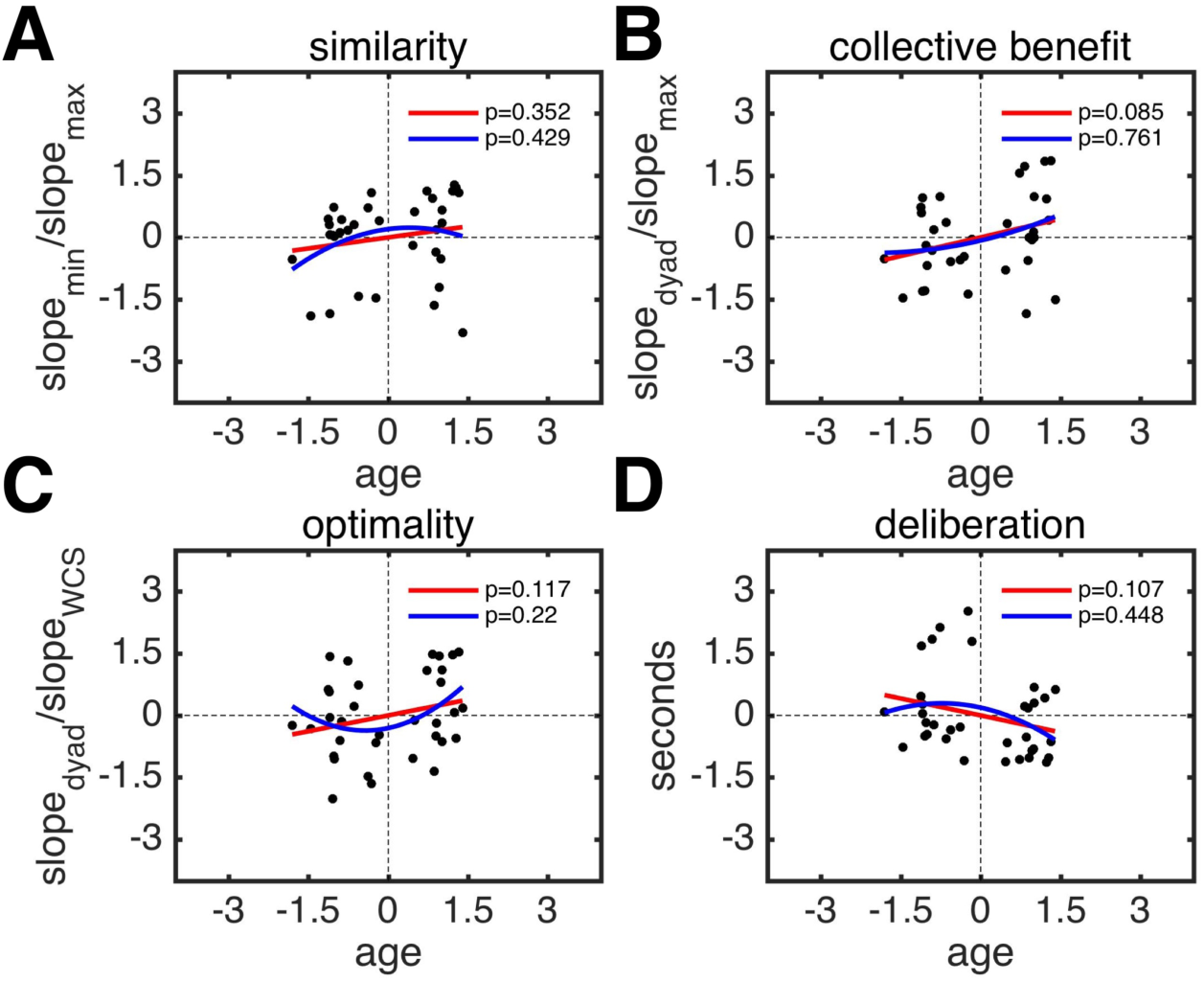
Developmental gradients for joint behaviour. **A**, Similarity of sensitivity. **B**, Collective benefit. **C**, Optimality. **D**, Deliberation time. **A-D**, Each dot represents a dyad. The lines indicate the slopes for the linear (red) and the quadratic (blue) terms for the mean age of dyads; the *p*-values indicate the significance of the terms. All variables were *z*-scored before being entered into the regression model.

